# The Inherited Rate Matrix algorithm for phylogenetic model selection for non-stationary Markov processes

**DOI:** 10.1101/2022.12.06.519392

**Authors:** Yurong Tang, Benjamin Kaehler, Hua Ying, Gavin Huttley

## Abstract

In phylogenetic reconstruction, substitution models that assume nucleotide frequencies do not change through time are in very widespread use. DNA sequences can exhibit marked compositional difference. Such compositional heterogeneity implies sequence divergence has arisen from substitution processes that differ between the lineages. Accordingly, there has been considerable interest in exploring the non-stationary model class, which is capable of generating divergent sequence composition. These models are typically parameter rich and this richness is further compounded by the use of time-heterogeneous variants in order to capture compositional heterogeneity. This illuminates two barriers limiting the widespread application of the non-stationary model class: both the compute-time required to fit a model and the risk of model over-fitting increase with the number of parameters and the extent of time-heterogeneity. In this study, we address these issues with a novel model selection algorithm we term the Inherited Rate Matrix algorithm (hereafter IRM). This approach is based on the notion that a species inherits the substitution tendencies of its ancestor. We further present the non-stationary heterogeneous across lineages model (hereafter ns-HAL algorithm) which extends the HAL algorithm of Jayaswal et al. [2014] to the general nucleotide Markov process. The IRM algorithm substantially reduces the complexity of identifying a sufficient solution to the problem of time-heterogeneous substitution processes across lineages. We also address the issue of reducing the computing time with development of a constrained-optimisation approach for the IRM algorithm (fast-IRM). Our algorithms are implemented in Python 3. From a simulation study based on 2nd codon position genome sequences of yeast, we establish that IRM is significantly more accurate than both ns-HAL and heterogeneity in the substitution process across lineages (hereafter HAL) for close and dispersed sequences. IRM is up to 70× faster than ns-HAL while fast-IRM is around 140× faster. fast-IRM also showed a marked speed improvement over a C++ implementation of HAL. Our comparison of the accuracy of IRM with fast-IRM showed no difference with identical inferences made for all data sets. These two algorithms greatly improve the compute time for model selection of a non-stationary process, increasing the suite of problems to which this important substitution model class can be applied.

## Introduction

Phylogenetic reconstruction methods are widely employed and the results are widely debated. A multitude of studies have shown that the results are sensitive to the taxa included [e.g. Rokas et al., 2003b,a, Reddy et al., 2017, Pisani et al., 2015] and that the tendency to generate conflicting trees is affected by choice of substitution model [e.g. Huttley, 2009]. Such discrepant results are likely to arise from model misspecification (incongruence between the assumptions made by the methods and the properties of the biological data to which they are applied). Biological sequence data arises by complex processes that act on species at the molecular and population level. The methodological component responsible for capturing that complexity is the substitution model and thus choice of suitable substitution model is crucial to the robustness of phylogenetic inference. Non-stationary substitution models are able to represent divergence in sequence composition. This class of models have been demonstrated to fit sequence data extremely well in comparison to the widely employed time-reversible substitution models [Kaehler et al., 2015]. The superior ability to describe the data of non-stationary models may cause over-fitting. [Kaehler et al., 2015] implemented bootstrap tests to efficiently identify over-fitting. In this work, we develop a new method aimed at reducing the risk of over-fitting.

At present, stationary substitution models are employed nearly exclusively. These models have the advantage of being relatively simple with a small number of parameters. However, they incorrectly assume that the composition of nucleotides, or amino acids, remains roughly the same across a tree [Felsenstein, 2003]. It has been demonstrated that violation of this assumption results in phylogenetic errors [e.g. Foster, 2004].

It has been known for decades that homologous DNA sequences can exhibit marked compositional difference in the relative abundance of the four nucleotides [Macaya et al., 1976, Jukes and Bhushan, 1986, Karlin and Mrázek, 1997]. Such observations indicate that the evolutionary process producing these sequences is not stationary; the sequence composition changes over time. This property has been the motivation for development of substitution models that can accommodate this non-stationarity. The mechanistic basis of this divergence in sequence composition can include contributions from both natural selection and changes in mutagenesis. For the sake of simplifying the arguments here, we consider only the influence of changes in mutation. It has been demonstrated that these models can give a markedly better description of sequence evolution than stationary ones [Kaehler et al., 2015]. However, a disadvantage of non-stationary models is they are very parameter rich compared to their time reversible counterparts.

Increasing the number of parameters can improve model fit. However, such increases in the number of parameters also makes a model prone to over-fitting [Burnham and Anderson, 2002] where the extra parameters fit noise in the data, inflating bias and causing errors in inference. In phylogenetic reconstruction, the assumption of heterogeneity in the substitution process across lineages is supported by evidence of compositional heterogeneity between the sequences [Jayaswal et al., 2014]. But an unrestricted accommodation of heterogeneity across lineages may contribute to overparameterising the model [Jayaswal et al., 2014]. A sensible reduction in parameter number can be achieved by restricting the number of rate matrices. However, the total number of possible ways of reducing heterogeneity among lineages is enormous for even a modest number of taxa [Jayaswal et al., 2011]. Therefore, an efficient strategy for model selection is required to identify an optimal time-heterogeneous model for a data set.

Multiple approaches for representing time-heterogeneous processes have been proposed. The most extreme possible choice is the most general model which allows a distinct non-stationary process for each edge of a phylogenetic tree [Jayaswal et al., 2011, Zou et al., 2012]. Another strategy is a distinct vector of equilibrium nucleotide frequencies on each edge of the tree [Yang and Roberts, 1995, Foster, 2004, Zou et al., 2012]. The former naturally represents non-stationary models but is more prone to over-fitting due to the larger number of free parameters. The latter is a composite of a time-reversible parameterisations with explicitly imposed shifts in the stationary distribution nodes of the tree, producing a non-stationary model (hereafter we term such models as pseudo non-stationary models). These models will also be susceptible to over-fitting.

One model complexity reducing approach employs a top-down algorithm to identify optimal and near-optimal time-heterogeneous models [Jayaswal et al., 2011]. The algorithm starts with the most general model (a distinct rate matrix on every edge) and then gradually decreases the model complexity until a stop condition is satisfied, producing a near-optimal model. This approach is only suitable for a small number of taxa. Subsequently, a bottom-up algorithm accounting for heterogeneous across lineages model (HAL) was developed [Jayaswal et al., 2014]. This algorithm starts with the simplest model and sequentially increases the model complexity until a stop condition is satisfied. In their approach, Jayaswal et al. [2014] employed a non-stationary model based on General Time Reversible rate matrices [Lanave and Saccone, 1984]. While the algorithm is comprehensive in terms of model reduction, it still has a massive solution space that makes its application impractical.

One notable attribute of the HAL algorithm is that disjoint edges can share the same rate matrix, which is distinct from the rate matrices operating on the intervening edges. This corresponds to a biological case of convergent evolution, whereby mutation and natural selection operate identically on edges that are not adjacent on the phylogenetic tree. While convergent evolution is plausible, it is a “special case” and not representative of the general tendencies of divergence between biological lineages.

Based on the limitations of the above approaches, it is apparent that new model selection strategies are required in order for non-stationary Markov processes to become a practical addition to the molecular evolutionary and phylogenetic toolkits. Systematic changes to mutation are important drivers for sequence substitution processes and changes to sequence composition [e.g. Karlin and Burge, 1995, Knight et al., 2001]. Mutation tendencies are determined, to a substantial degree, by DNA modification and repair systems which are transmitted from parent to offspring. Accordingly, it seems reasonable to conjecture that lineages inherent their mutation tendencies (and thus substitution processes) from their immediate ancestor. Thus, lineages whose immediate ancestor is the same are most likely to exhibit the same substitution process. For instance, gene loss(es) in some lineages have led to complete absence of the hypermutable base 5-methyl-cytosine [Albalat et al., 2012]. Descendants of those lineages thus exhibit a significant shift in the relative abundance of the different point mutations. Motivated by this conjecture, we propose a bottom-up Inherited Rate Matrix (IRM) algorithm for selection of time-heterogeneous non-stationary models.

Here we report our evaluation of the question of whether a model selection strategy based on inheritance of rate matrices can prove sufficient in model selection of non-stationary models. IRM algorithm coupled with an efficient tree traversal mechanism are described. We further present a non-stationary HAL algorithm (ns-HAL) based on the non-stationary general Markov nucleotide substitution model [Kaehler et al., 2015]. We show via simulation that IRM is fast and less prone to model misspecification than ns-HAL. The source codes of the methods presented in this paper are made available at http://doi.org/10.5281/zenodo.3754681. The raw data, results and scripts used for analyses are available at http://doi.org/10.5281/zenodo.3754674.

## Materials and Methods

### Data used in this study

We used the same data set as that employed by [Jayaswal et al., 2014]. This data set consists of the concatenation of second codon positions from 106 multiple sequence alignments of orthologs from eight yeast species. The resulting concatenated alignment is 42,337bp long. The species, and their corresponding abbreviation are: *Saccharomyces cerevisiae* (Scer), *Saccharomyces paradoxus* (Spar), *Saccharomyces mikatae* (Smik), *Saccharomyces kudriavzevii* (Skud), *Saccharomyces castellii* (Scas), *Saccharomyces kluyveri* (Sklu), *Saccharomyces bayanus* (Sbay), and *Candida albicans* (Calb). A phylogenetic tree for these species reported by of Rokas et al. [2003b] is shown in Figure 1. The alignment, a concatenation of alignments of one-to-one orthologs determined by [Rokas et al., 2003b], was employed by [Jayaswal et al., 2014] in development of the HAL algorithm.

**Figure 1:**
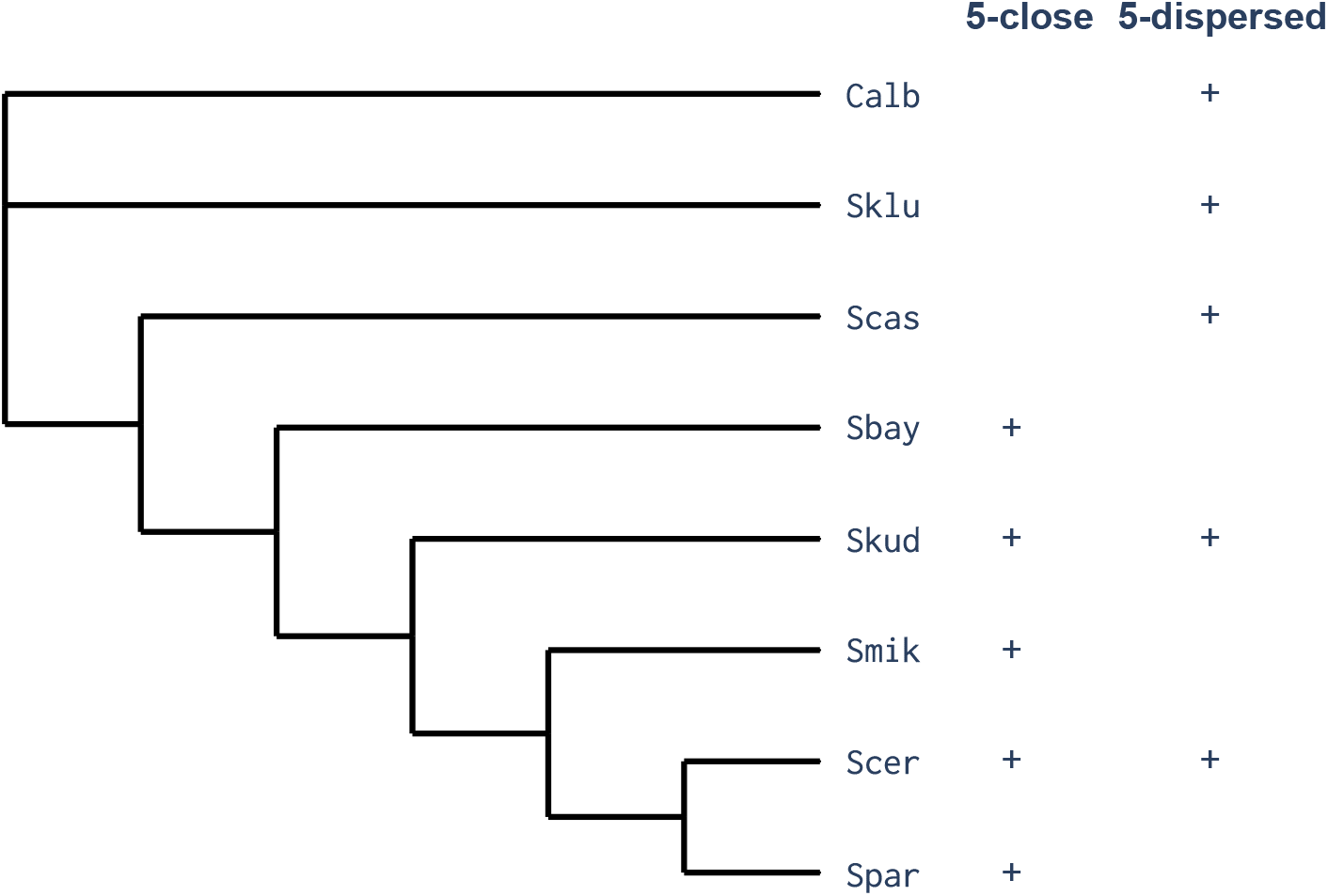
The phylogeny of the 8 Yeast species from Rokas et al. [2003b]. The eight species are *S. cerevisiae* (Scer), *S. paradoxus* (Spar), *S. mikatae* (Smik), *S. kudriavzevii* (Skud), *S. castellii* (Scas), *S. kluyveri* (Sklu), *S. bayanus* (Sbay), and *C. albicans* (Calb). Membership of the 5-close species subset {Scer, Spar, Smik, Skud, Sbay} and 5-dispersed species subset {Scer, Skud, Scas, Sklu, Calb} are shown in the corresponding columns using the ‘+’ symbol.

In order to decrease running time of ns-HAL, we selected two 5-species subsets from the data. The “5-close” subset consists of the five most closely related lineages Scer, Spar, Smik, Skud and Sbay. The “5-dispersed” subset consists of five more dispersed lineages Scer, Skud, Scas, Sklu, Calb. The topologies of 5-close data set and 5-dispersed data set are subgraphs from the topology of Figure 1. We expected that the extent of divergence between a collection of taxa would influence power to resolve different levels of time-heterogeneity. The 5-close and 5-dispersed subsets were chosen to facilitate the examination of this hypothesis. How this was done is explained in the section on statistical evaluation.

### General Markov nucleotide substitution model

The first non-stationary general Markov nucleotide substitution model to be studied was that of [Barry and Hartigan, 1987] (hereafter referred to as the BH model). This is a discrete-time, non-stationary Markov process for nucleotides that has 12 different parameters. The most general continuous-time Markov process on nucleotides [hereafter GN Kaehler et al., 2015] is equivalent to BH if the Markov process is embeddable [Verbyla et al., 2013]. In GN, all possible exchanges receive their own rate parameter, thus there are 12. In Cogent3 [Huttley, 2019], the rate matrix is calibrated such that −Σ_*i*_ *π*_*i*_ *Q*_*ii*_ = 1. For a stationary process, this is the expected number of substitutions in a unit time interval and, as a result, the scalars applied per branch are exactly the expected number of substitutions per site for that branch. For a non-stationary process, *π* changes through time and thus that relationship no longer holds. This choice of calibration is a convenience for the purpose of most model fitting and the correct expected number of substitutions for each branch can be derived after fitting [Kaehler et al., 2015]. For the GN model developed by Kaehler et al. [2015], the substitution number, genetic distance and ENS distance are the same. Therefore, the above three terms described in this paper are the same.

Sufficient conditions under which the BH model is identifiable were established by [Chang, 1996] and we refer the reader to that article for a complete exposition of these properties. One sufficient condition is that rate matrices exhibit the property of diagonal largest in column (hereafter DLC). Identifiability of the continuous-time general Markov nucleotide counterpart to BH [Kaehler et al., 2015] requires in addition that there is a unique mapping between *P* (*t*) = exp^*Qt*^ [Kaehler et al., 2015]. Hence, we employed two identifiability checks: (i) the estimated processes exhibit DLC, and (ii) a unique mapping exists [Kaehler et al., 2015]. Solutions from numerical optimisation that failed these checks were discarded.

### Identifying the direction of “time”

As shown by [Chang, 1996], the general Markov process cannot identify the direction of time. Stated another way, such a process is not identifiable for an unrooted tree of three taxa. This makes it problematic to identify descendants from an edge, a key prerequisite for IRM and ns-HAL. Of the possible approaches to solve this, we used the midpoint method as it is computationally efficient (needs to be done only once) and was an existing Cogent3 tree capability. In essence, we orient the tree such that the “root” node is as close to being the chronological root as possible. Once this is done, the assumption that children of a node are its actual descendants seems reasonable.

### ns-HAL: HAL algorithm for non-stationary Markov processes

In our ns-HAL algorithm, we made the following modifications to the HAL algorithm of Jayaswal et al. [2014]:

- *Use the GN model* Jayaswal et al. [2014] employed a non-stationary model based on the time-reversible GTR process. It achieved non-stationarity by associating distinct GTR matrices with different stationary frequency vectors (*π*). We do not need to do this as the general Markov nucleotide (GN) process is naturally non-stationary [Kaehler et al., 2015].
- *Use an unrooted tree* Jayaswal et al. [2014] specified HAL for a strictly binary tree, hence its root node has two branches. As a GN model with a rooted tree is not identifiable, we used an unrooted tree.
- *Decrease model complexity using Frobenius norm [Golub and Van Loan, 2013, p. 70]* In order to avoid overparameterization, Jayaswal et al. [2014] applied a method of decreasing model complexity [the CORE algorithm, Jayaswal et al., 2011] to determine whether the first model in set S produced by HAL is overparameterized. In their method, clustering of *π* was used to guide model complexity reduction. As GN does not require distinct *π* for a time-heterogeneous non-stationary process, we used the Frobenius norm between the two closest rate matrices, combining them into one rate matrix so that a simpler model is obtained.
- *Different model selection criteria* Jayaswal et al. [2014] used Akaike information criteria (*AIC*) and Bayesian information criterion (*BIC*) for selecting models. In addition to the above two information criteria, we also evaluated using *AIC* correction (*AICc*) for model selection as it has a stronger penalty for the number of parameters than AIC.
- *Different iteration stop condition* In Jayaswal et al. [2014], the iteration stop condition is controlled by the size of a set S. In each iteration, a fixed number of candidate models are put into this set. In ns-HAL, candidate models can be selected through two methods: (i) use a fixed number of candidate models as done by Jayaswal et al. [2014], (ii) choose candidate models according to *AICc* difference.
- *Identifiability check* Tests of the two identifiability conditions, DLC and Unique mapping [Kaehler et al., 2015] are performed to check if the rate matrix is identifiable. Jayaswal et al. [2014] do not perform this step.

### IRM: Inherited Rate Matrix algorithm for non-stationary Markov processes

Motivated by the greater likelihood of lineages to inherent the substitution properties of their immediate ancestor, we impose the restriction that only adjacent edges can share a rate matrix. A phylogenetic tree is an acyclic connected graph in which any two nodes are connected by exactly one path constituted by some joint edges. In order to guarantee that all edge set partitions consist of adjacent edges, we adapted a graph cut algorithm.

In graph cut theory, a cut is a partition of the nodes (vertices) of a graph into two disjoint subsets. An important and relevant partition algorithm is that of minimum cut. The minimum cut algorithm partitions nodes of graph into two sets according to the weight of edges, choosing the edge with the smallest weight. Our application here is different, as we seek to obtain a partition of the edges of a graph instead of a partition of the nodes. Therefore, we need to cut edges into two groups from one node (we denote this node as the cutting node). The notion of a weight is used to determine the order of cutting nodes.

We developed a variant algorithm of minimum cut, which is illustrated in Figure 2. In it, we choose cutting nodes according to weights of branches and then partition edges into two groups. We define the weight as 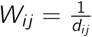, where *d*_*ij*_ is the genetic distance from the node *i* to the node *j*. We choose the cutting node by a genetic distance based weight for the following reasons. The genetic distance is a quantity derived explicitly from the rate matrix. It is related to the amount of information in the data regarding a specific edge. Finally, the probability of a change in substitution process, and hence the rate matrix, increases with increasing chronological time. Our choice of edge weight should therefore identify the edge that is most likely to have evolved in a manner described by a distinct rate matrix.

**Figure 2:**
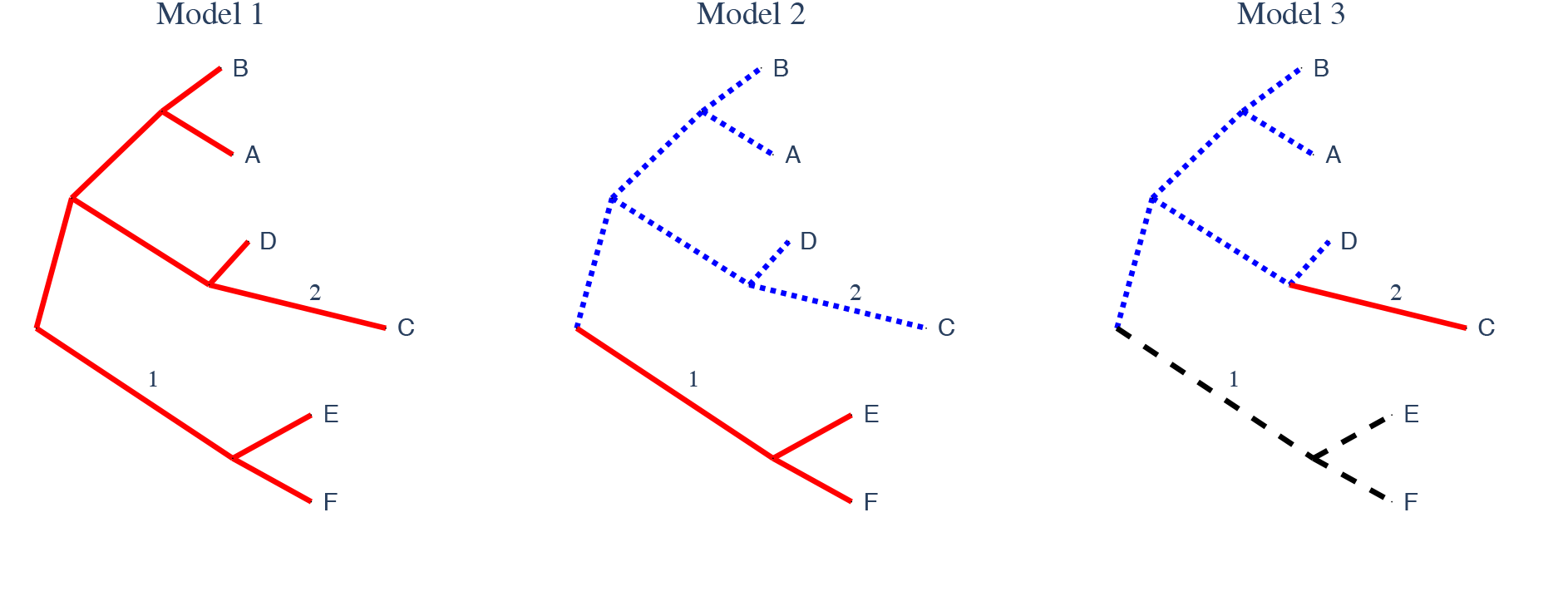
An example of the IRM and fast-IRM algorithms. Model 1 is the parent model of Model 2 which is the parent of Model 3. In each panel, edges with the same colour share a rate matrix. There is one rate matrix for Model 1, 2 rate matrices for Model 2 and 3 rate matrices for Model 3. The algorithm compares the fit of Model 2 to that of Model 1 using a nominated statistic like AICc. It accepts a model if that fit exceeds a minimum threshold and proceeds to consider the next model. fast-IRM is distinguished from IRM by making the parameters from the parent model constant to reduce numerical optimisation time. For instance, in Model 3, the edges coloured black are constrained to be constant at the estimated values of the parent (Model 2). All other parameters are free to be optimised.

The implementation of our variant minimum algorithm includes two steps. First, with the edges ranked in ascending order by their weight, an iterative process is then applied. Parent node of the edges having the smallest weight is then chosen as the cutting node. The cutting node defines two sets of edges. Edges within a set share the same rate matrix. (Note that all edges retain an independent scalar parameter that is monotonic to genetic distance [Kaehler et al., 2015].) In cases where there are ties in weight, all edges with the equivalent weight are evaluated. Second, an iteration stop condition is evaluated for every new choice of cutting node. An information theoretic measure, such as *AIC*, is employed. Simplified Python implementations of the core IRM algorithm and associated utility functions are presented in Supplementary Materials Algorithm S1-S3.

### fast-IRM: IRM with constrained optimisation

The computational speed of algorithms for model selection depends on whether full numerical optimisation is used for evaluating each candidate model. In a full numerical optimisation, all free parameters are subject to modification during the optimisation. For a general nucleotide Markov model, this corresponds to 11 free parameters per rate matrix and a unique branch scalar for each edge that is monotonic to the expected number of substitutions [Kaehler et al., 2015]. In both the IRM and ns-HAL cases, maximum likelihood estimates from the parent (simpler) model were used as initial values for each derived model. Here, a derived model was generated from a parent model according to the model selection algorithms. In order to further speed the algorithm, we developed a constrained-optimisation variant of IRM that we denote fast-IRM.

In fast-IRM reduces compute time by decreasing the number of parameters optimised at each step. When a clade is split into two at the cutting node, the two resulting clades are represented by 2 rate matrices. In IRM, the entire model is numerically optimised. Hence, the number of free parameters to be numerically optimised increases with each iteration. In fast-IRM, only parameters relating to the 2 new rate matrices are optimised whilst all other parameters are fixed as constant at the parent model optimised values. For instance, the cutting node that defines Model 3 in Figure 2 splits the blue clade of Model 2. The black edges in Model 3 lie outside of this and thus their parameters are fixed constant at the values from the red clade of Model 2 (black edges in Figure 2). The number of free parameters at each iteration of fast-IRM is therefore fixed as 2× the number per rate matrix plus the total number of edges in the clade being split (6 in Model 3, Figure 2).

For the GN model, where each additional rate matrix introduces 11 new rate terms, the numerical optimisation performed at each iteration through fast-IRM is limited to 22 rate terms plus a branch scalar per edge within the clade. Thus, for Model 3 in Figure 2, there are 28 free parameters. The fast-IRM algorithm is an approximate likelihood method and thus the convergence property of maximum likelihood is not guaranteed.

### Size of the solution space

We contrast the solution space size (number of possible models) for 5-taxa for ns-HAL, IRM and the theoretical maximum. For the two algorithms, we determined the maximum number by exhaustive evaluation using the implemented algorithms. The theoretical maximum is a Bell number [Jayaswal et al., 2014]. The solution space size for differing values of *k* is shown in Table 1. A *k* = 1 corresponds to a model in which there is a single rate matrix for which there is only one possible model. When *k* = 7, every edge has a separate rate matrix for which there is only one possible model. In these two extreme cases, all algorithms have the same solution space size. For *k* ∈ {2, …, 6} the number of possible solutions for ns-HAL grows substantially due to possible ways of combining edges. In contrast, because the IRM algorithm uses mincut only a single additional edge can be considered at each iteration through possible values of *k*. The exception occurs if there are tied branch length values, in which case the solution space for IRM will equal the number of tied values.

**Table 1:** Comparison of the number of possible models to be evaluated for a 5-taxa tree. *k* is the number of rate matrices, Maximum is the maximum size of the solution space (a Bell number) for *k*, numbers under the IRM and ns-HAL columns correspond to the solution size determined by exhaustive search using our implementation of the algorithms.

| **k** | **Maximum** | **ns-HAL** | **IRM** |
| --- | --- | --- | --- |
| 1 | 1 | 1 | 1 |
| 2 | 63 | 9 | 1 |
| 3 | 301 | 38 | 1 |
| 4 | 350 | 75 | 1 |
| 5 | 140 | 50 | 1 |
| 6 | 21 | 13 | 1 |
| 7 | 1 | 1 | 1 |

### Statistical evaluation of IRM, ns-HAL and HAL

We calculated the accuracy with which the IRM, ns-HAL and HAL algorithms correctly identified the generating model. We indicate the model used to generate the simulated alignments as the *true model*. Specifically, the true model is characterised by the set of *k* distinct rate matrices and the *k* edge sets, denoted 𝔼_*k*_, to which they map. The model with *n* rate matrices and the edges to which they map (i.e. 𝔼_*n*_) identified by an algorithm on a simulated alignment is referred to as the *inferred model*, or *estimated model*. We consider an inferred model correct when 𝔼_*k*_ = 𝔼_*n*_. In words, the inferred model was correct if it found the same partition of the tree into edge sets as that used to simulate the alignment. The relative performance of each algorithm was established by comparing the total count, obtained across all model conditions, of correctly inferred models.

Simulated alignments were generated based on a fit of the GN model to the yeast data. The specification of time-heterogeneity was done in a manner consistent with the assumption of rate matrix inheritance. The model with a single rate matrix is the first fitted model. Then the second fitted model was generated where the edges descending from one edge with the biggest genetic distance has one rate matrix where all the other edges have a different rate matrix. The process, choosing an edge according to genetic distance order and a distinct rate matrix is assigned to the edges descending to the chosen edge, continue until the last model with the maximum number rate matrices (*k*) was generated. A 5-species tree has 7 edges so the maximum number of *k* is 7. For each value of *k*, all possible time-heterogeneous models consistent with IRM were fit to both the 5-close and 5-dispersed yeast data sets. From each such fitted model, 10 simulated alignments with the same length with fitted model were generated. The IRM, ns-HAL and HAL algorithms were then applied to each generated model.

As each algorithm was applied to the same simulated alignments, we have a paired study design. We illustrate how algorithms were compared using IRM and ns-HAL. The paired design allows producing a 2 × 2 contingency table with columns corresponding to whether the IRM algorithm correctly recovered the generating model, or not. The rows correspond to the equivalent outcome for the ns-HAL algorithm. A McNemar test [Edwards, 1948, Roggo et al., 2003], with correction for small samples, was then used to test the null hypothesis that the performance of two algorithms was the same.

### Software implementation

The algorithms described above were implemented in Python-3.7 or greater and Numpy 1.18.1. Core functions for likelihood calculation and numerical optimisation are provided by Cogent3 [Huttley, 2019]. Implementation accuracy was validated using unit tests and these tests covered ∼86% of the code [Batchelder, 2019]. Version 1.09 of HAS-HAL, written in C++ [Jayaswal et al., 2014, Wong, 2017] was used. The source codes of the methods presented in this paper are made available at http://doi.org/10.5281/zenodo.3754681. The raw data, results and scripts used for analyses are available at http://doi.org/10.5281/zenodo.3754674.

## Results

We briefly reiterate the experimental design here. We denote the number of rate matrices in a phylogenetic model as *k*. For a phylogenetic tree of 5 taxa, and 7 edges, *k* can be at minimum 1, and at maximum 7. We generated synthetic data for each possible value of *k* for the 5-close and 5-dispersed yeast data sets. These data were used to contrast the accuracy of the algorithms, their computational speed and the consistency between the IRM and fast-IRM variants.

### IRM is more accurate than ns-HAL and HAL

For a specific simulated alignment, we considered an algorithm accurate if it correctly recovered the configuration of rate matrices corresponding to the generating model (i.e. 𝔼_*k*_ = 𝔼_*n*_). For assessing the accuracy calculation of HAL, we used *AIC* information criteria based on the same simulated alignments used by ns-HAL and IRM. Table 2 indicates the accuracy on the 5-close and 5-dispersed species data, respectively. The accuracy of HAL is is better than ns-HAL when *k* = 2, *k* = 5 and *k* = 6 for 5-close. However, the accuracy of ns-HAL is better than HAL when *k* = 1 and *k* = 3. In total, ns-HAL have nearly the same accuracy with HAL for 5-close. For 5-dispersed, ns-HAL is better than HAL for all the *k*. IRM exhibited higher accuracy than both HAL and ns-HAL when *k* is 1, 2, 3, 4 and 7 on 5-close species. For *k* values from 5 to 6, IRM was at least as good as ns-HAL. The three algorithms had low accuracy for *k* vales from 4 to 7 for 5-close. There was a tendency on the 5-close data set (which has more closely related sequences) for the algorithms to identify optimal models with few rate matrices. For 5-dispersed species, IRM had systematically higher accuracy than both HAL and ns-HAL.

**Table 2:**
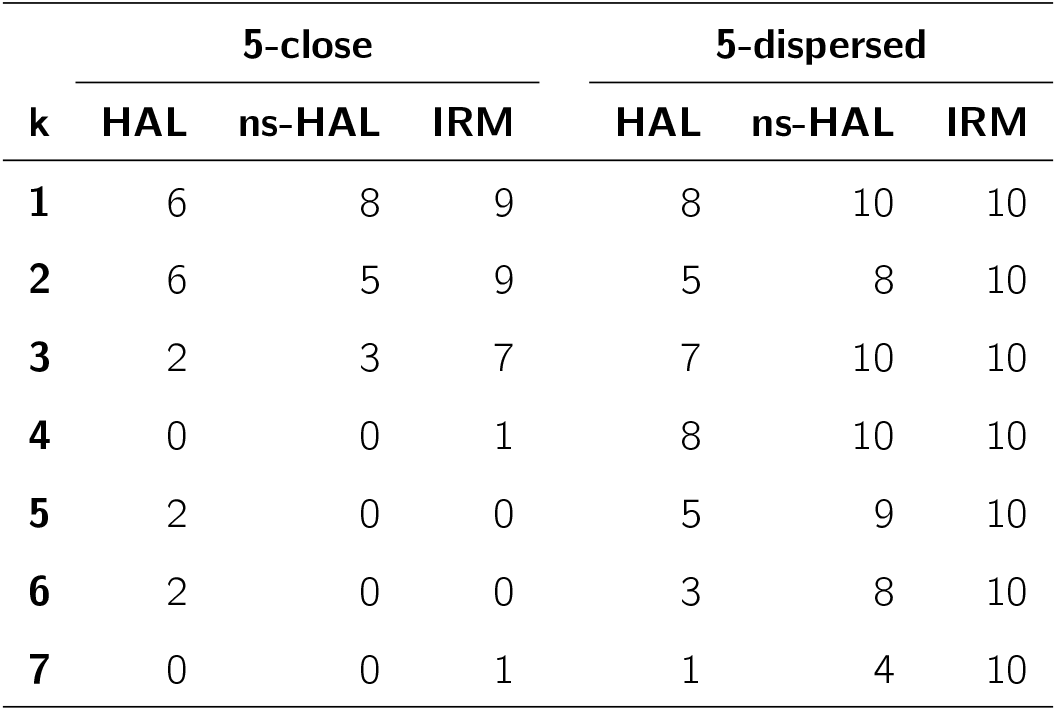
Accuracy comparison between the IRM, ns-HAL and HAL algorithms. 5-close and 5-dispersed are the different data sets. *k* is the number of rate matrices used to generate the 10 replicate data sets. Numbers correspond to the counts of when the inferred models returned respectively by IRM, ns-HAL and HAL were correct.

| **k** | **5-close** |  |  | **5-dispersed** |  |  |
| --- | --- | --- | --- | --- | --- | --- |
|  | **HAL** | **ns-HAL** | **IRM** | **HAL** | **ns-HAL** | **IRM** |
| **1** | 6 | 8 | 9 | 8 | 10 | 10 |
| **2** | 6 | 5 | 9 | 5 | 8 | 10 |
| **3** | 2 | 3 | 7 | 7 | 10 | 10 |
| **4** | 0 | 0 | 1 | 8 | 10 | 10 |
| **5** | 2 | 0 | 0 | 5 | 9 | 10 |
| **6** | 2 | 0 | 0 | 3 | 8 | 10 |
| **7** | 0 | 0 | 1 | 1 | 4 | 10 |

In order to verify if IRM is significantly different to ns-HAL, we applied the continuity corrected version of the McNemar test. Two 2 × 2 contingency tables were generated on the 5-close data set and 5-dispersed data set (Table 3). These contrast the ability of the two algorithms to recover the correct model. For both the 5-close and 5-dispersed data sets, we rejected the null hypothesis that the two algorithms were equivalent with p-values of ∼0.0026. Given its systematically higher accuracy (Table 2), we conclude that IRM is significantly more accurate than ns-HAL. Our comparison of ns-HAL and HAL (Table 4) indicated that ns-HAL is as accurate as HAL for 5-close and is significantly more accurate than HAL for 5-dispersed.

**Table 3:** Comparison of algorithm accuracy for IRM and ns-HAL. As each algorithm was applied to exactly the same simulated data we have a paired study design. From the 70 simulated alignments we record in the 2 × 2 table how often the algorithms were in agreement. Here, “True” means the algorithm recovered the correct model, “False” indicates it did not. Using the McNemar test with correction, we reject the null hypothesis of no difference between the algorithms. The p-values for both 5-close and 5-dispersed were ∼ 0.0026.

| **IRM** |  |  |  |  | **IRM** |  |  |  |  |
| --- | --- | --- | --- | --- | --- | --- | --- | --- | --- |
|  |  | **True** | **False** | **Total** |  |  | **True** | **False** | **Total** |
| **ns-HAL** | **True** | 16 | 0 | 16 | **ns-HAL** | **True** | 59 | 0 | 59 |
|  | **False** | 11 | 43 | 54 |  | **False** | 11 | 0 | 11 |
|  | **Total** | 27 | 43 | 70 |  | **Total** | 70 | 0 | 70 |

(a) 5-close
(b) 5-dispersed

**Table 4:**
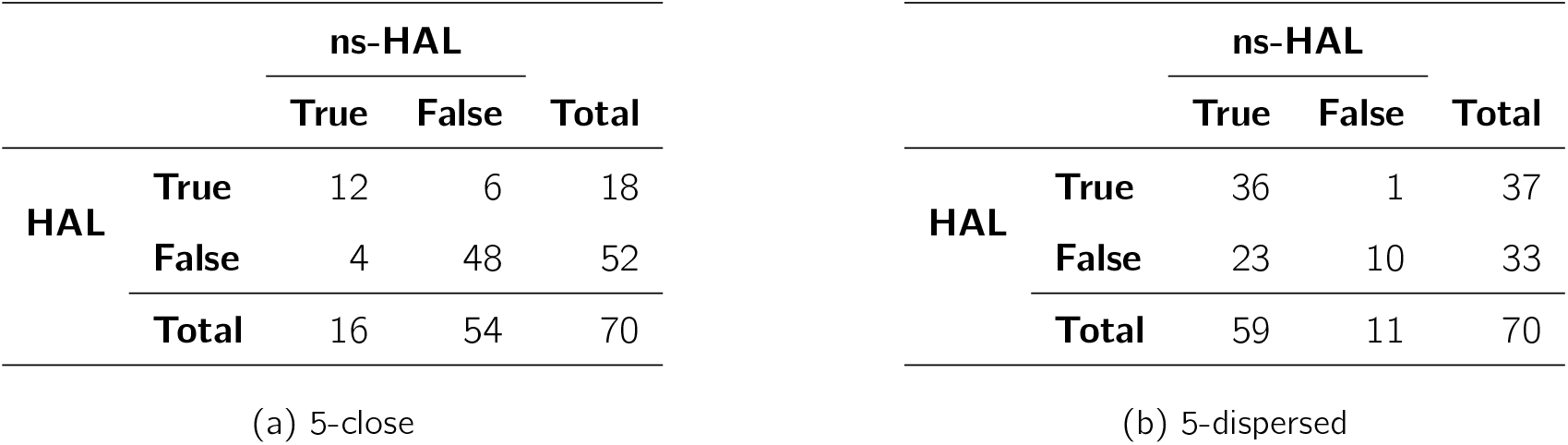
Comparison of algorithm accuracy for ns-HAL and HAL. As each algorithm was applied to exactly the same simulated data we have a paired study design. From the 70 simulated alignments we record in the 2 × 2 table how often the algorithms were in agreement. Here, “True” means the algorithm recovered the correct model, “False” indicates it did not. Using the McNemar test with correction, we reject the null hypothesis of no difference between the algorithms. p-values were ∼ 0.75 and ∼ 1.81*e* − 05 for 5-close and 5-dispersed, respectively.

We evaluated the potential contribution of differing statistical power to the difference of accuracy between 5-close and 5-dispersed. The relationship among the 5-close taxa indicates there will be less genetic distance separating these sequences. As the model is being fit to substitution events, a smaller number of events may impact on resolution. We determined the difference between the two data sets for the two algorithms in the estimated total genetic distance. As shown in Supplementary Materials Table S2, there is roughly an order of magnitude more events in the 5-dispersed data compared with 5-close. The likely impact of such a difference on statistical power seems the most plausible explanation for the difference accuracy between the data sets.

### Accuracy of fast-IRM

In order to assess the efficacy of the fast-IRM constrained-optimisation approach we contrasted its accuracy with that of IRM. In our implementation, it is in fact full optimisation when *k* is 1 and 2. Therefore, we only compared the accuracy when *k* ∈ {3, 4, … 7}. Table 5 shows that constrained-optimisation and full-optimisation have the same accuracy for the 5-dispersed species when *k* ranges from 3 to 7 and near identical accuracy for the 5-close species. The 2 × 2 contingency tables (Table 6) from the 5-close data set and 5-dispersed data set, revealed the algorithms recovered identical edge sets on each alignment with the single exception of 5-close where *k* = 6.

**Table 5:** Accuracy comparison between algorithm IRM and algorithm fast-IRM. Here, *k* is the rate matrix number. The numbers of the column “IRM” and the column “fast-IRM” are correct number of inferred models returned respectively by IRM and fast-IRM based on 10 simulated alignments.

| k | 5-close |  | 5-dispersed |  |
| --- | --- | --- | --- | --- |
|  | IRM | fast-IRM | IRM | fast-IRM |
| **3** | 7 | 7 | 10 | 10 |
| **4** | 1 | 1 | 10 | 10 |
| **5** | 0 | 0 | 10 | 10 |
| **6** | 0 | 1 | 10 | 10 |
| **7** | 1 | 1 | 10 | 10 |

**Table 6:**
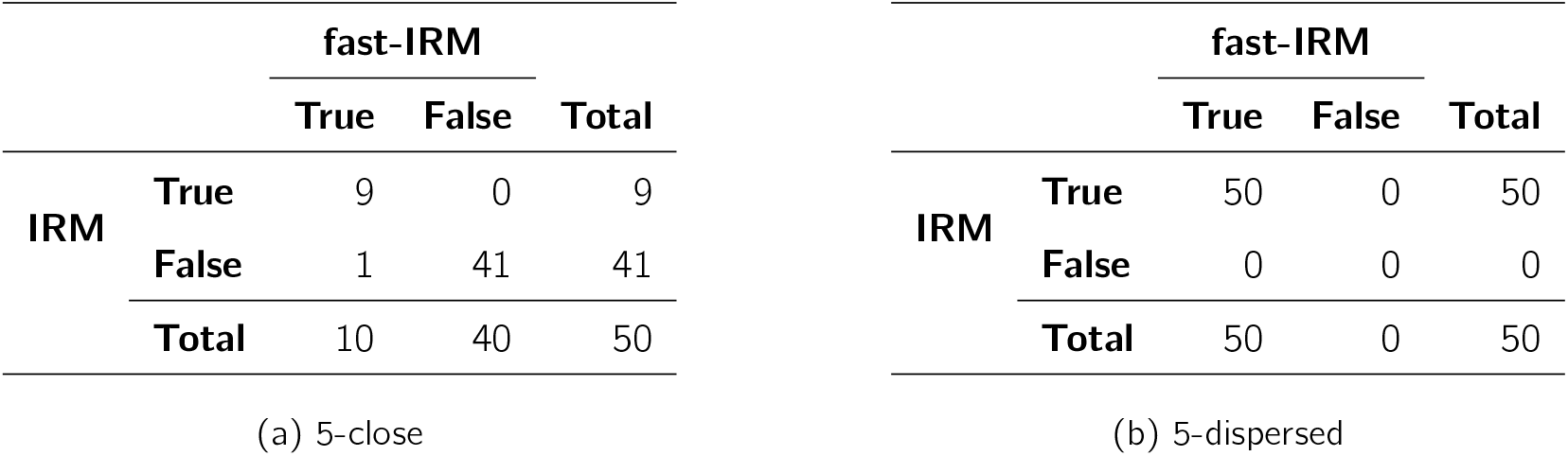
Inferences from IRM and fast-IRM were almost perfectly congruent. From the 50 simulated alignments where *k* ∈ {3, 4, … 7}, we record in the 2 × 2 table how often the algorithms were in agreement.

We assessed whether the fast-IRM algorithm resulted in log-likelihoods that were different to those from IRM. The differences in log-likelihood for each value of *k* are presented in Figure 3. The absolute difference in the log-likelihood is tiny when *k* ∈ {3, 4, … 7} for 5-close species. While there were some outlier points, the absolute value of those difference was big for *k* = 6. For 5-dispersed species, however, the differences were larger with an Interquartile Range (IQR) ranges of around 4. As *k* increased, the difference increased also. Despite these discrepancies, the choice of optimal model was congruent between the two algorithms.

**Figure 3:**
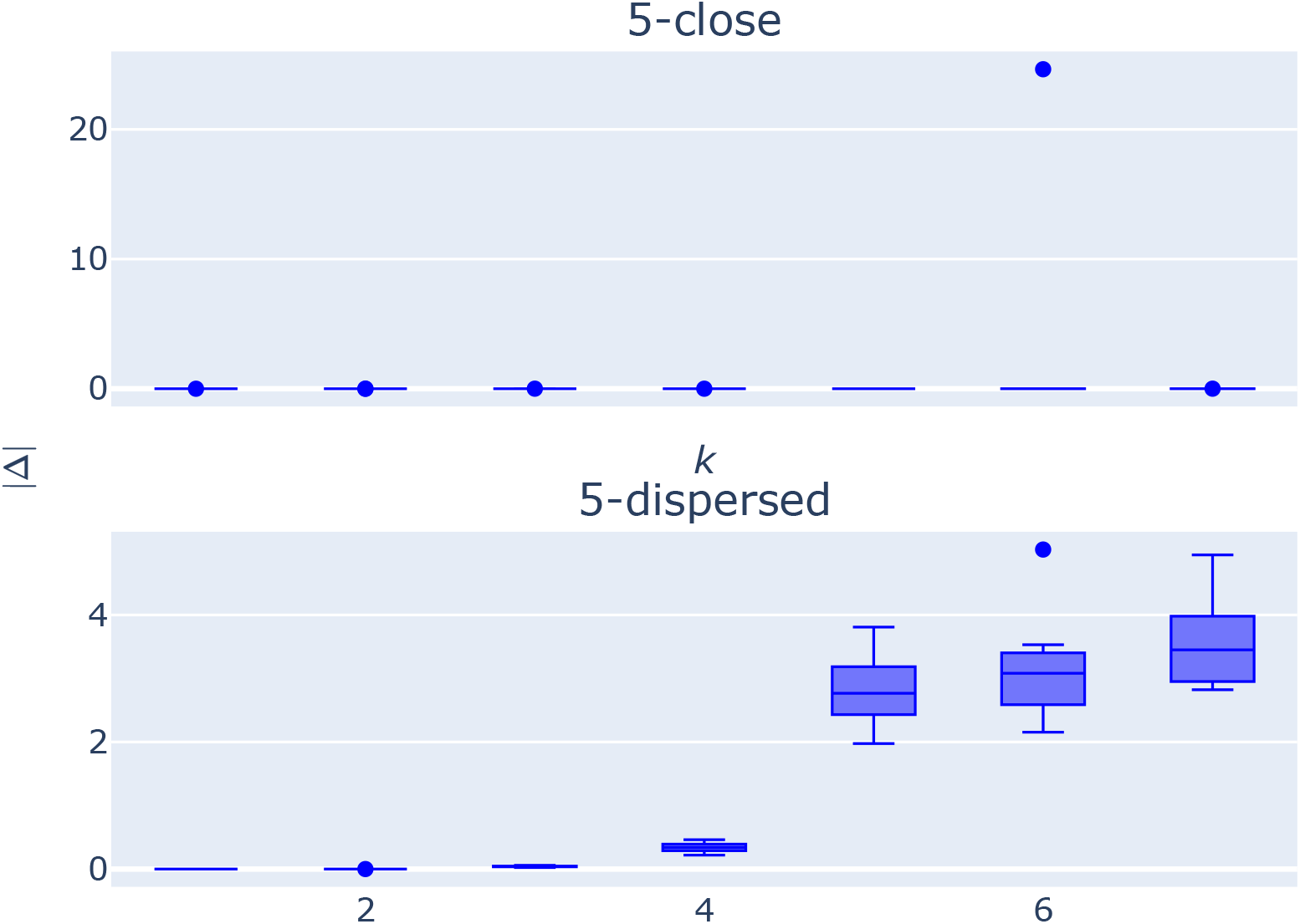
Consistency in log-likelihood between the IRM and fast-IRM algorithms. The y-axis is | Λ |=| log *L*(IRM) − log *L*(fast-IRM) |. The x-axis is *k*, which is the number of distinct rate matrices.

### Computational performance

We compared computational time required to identify the optimal model between ns-HAL, IRM and fast-IRM. The comparison was fair in so much as performance was measured from evaluation of the same simulated data sets as those used for assessing statistical performance. The three algorithms were run on a shared high-performance computing environment. Each simulated alignment of each *k* was assigned to one compute node of this environment. Each node had 2× 8 core Intel Xeon E5-2670 (Sandy Bridge) 2.6GHz. Then the median and quartiles of running times from 10 simulated alignments for each *k* are displayed (Figure 4).

**Figure 4:**
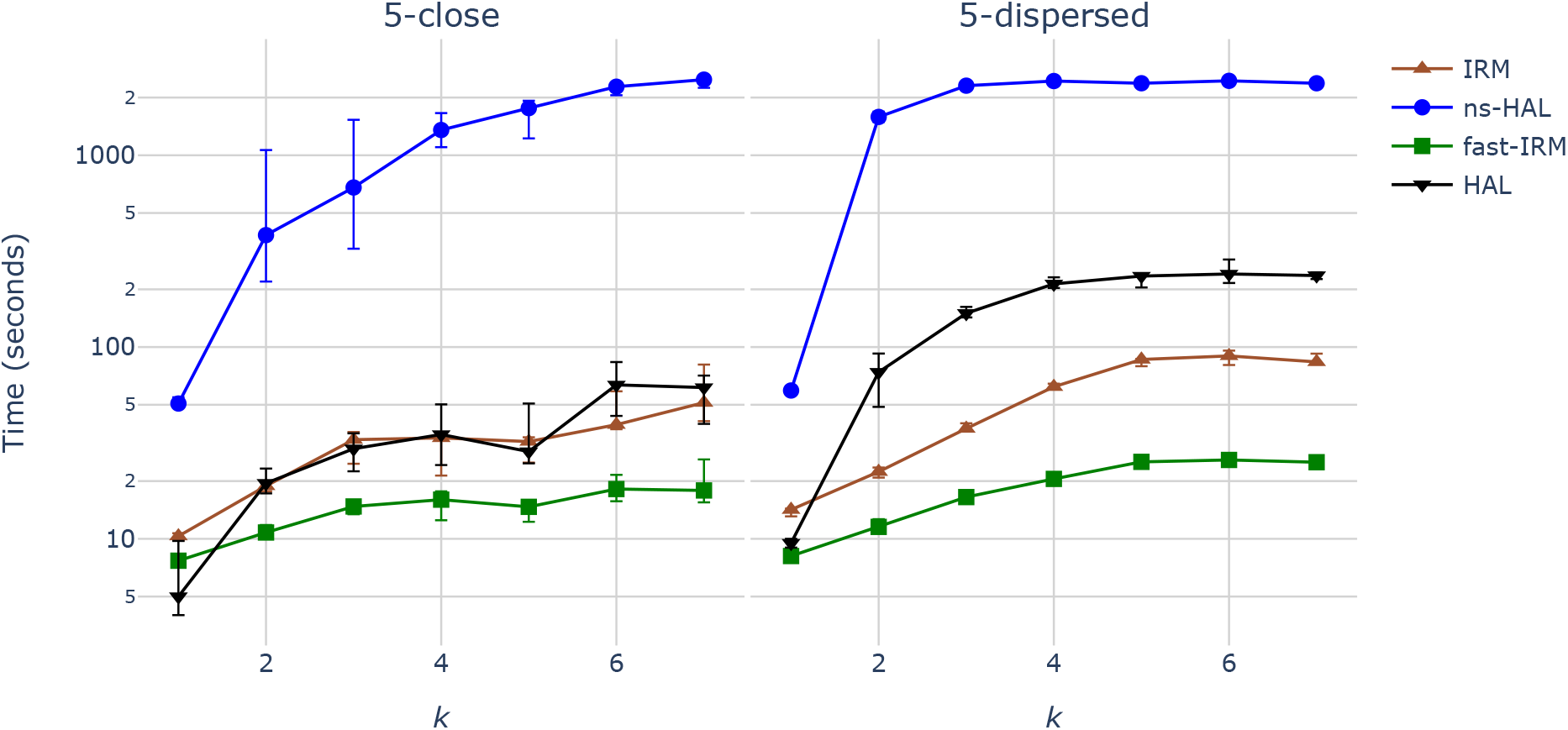
Median running time (*log*_10_-scale) of HAL, ns-HAL, IRM and fast-IRM. The data set is indicated by the subplot title. Rate matrix number, *k*, is shown on the x-axis. The y-axis is the median of log_10_(*seconds*) across the 10 replicates and the error bars being the lower and upper quartiles.

Our results established that both variants of IRM were markedly faster than ns-HAL. The compute time of ns-HAL ranged from ∼ 8× (*k* = 1) to ∼ 30× (*k* = 2) slower than IRM for 5-close species. For the 5-dispersed data set, the relative speed ranged from ∼ 5× (*k* = 1) to ∼ 75× (*k* = 2) slower than IRM. The speed relative to fast-IRM was considerably worse, with ns-HAL speeds slower by as much as ∼ 200× (*k* = 2). The slowdown from using IRM compared to fast-IRM ranged from ∼ 1.6× (*k* = 1, 5-close) to ∼ 3.4× (*k* = 7, 5-dispersed). fast-IRM was systematically faster than HAL for both 5-close and 5-dispersed data sets (with the exception of *k* = 1). The speed of IRM was faster than HAL except for *k* = 1 and *k* = 7 for 5-close and except *k* = 1 for 5-dispersed.

### Consistency between IRM and fast-IRM

On 50-taxon empirical coral data with the second codon sites, on NCI, we compared the optimal models obtained by IRM and fast-IRM and their running time shown in the Table 7. We found that their optimal models are exactly the same and their rate matrix number is 4.

**Table 7:** Optimal model number and running time on naked coral data. *k* is the number of rate matrices and *t* is the running time

|  | IRM | fast-IRM |
| --- | --- | --- |
| k | 4 | 4 |
| t(sec) | 175.0903 | 80.7143 |

## Discussion

We developed two algorithms, IRM and ns-HAL, for selection of the optimal non-stationary model for a given tree topology. In essence, both algorithms are bottom-up approaches; they start with the simplest model (one rate matrix) and gradually increase model complexity (number of rate matrices) of the model until a stop condition is satisfied. For a tree with large taxa number, bottom-up approaches should be faster than top-down approaches because the latter start with the most complex (parameter rich) model. When the number of taxa is large, the total solution space dramatically increases. Our results indicated that IRM or fast-IRM should be preferred over ns-HAL or HAL because of greater accuracy and markedly less compute-time.

Our comparison of algorithm accuracy established IRM was better than ns-HAL (Table 2). For 5-close species, IRM had high accuracy for *k* ∈ {1, 2, 3} but ns-HAL had high accuracy only for *k* = 1. In contrast, both algorithms showed systematically greater accuracy for 5-dispersed species compared to their 5-close results. IRM produced perfect accuracy for all values of *k* while ns-HAL showed a sharp reduction in accuracy for larger *k* on 5-dispersed data and it generated same or higher accuracy for each *k* than ns-HAL on 5-close (Table 2). The differences between the two data sets likely derives from differences in statistical power. The total number of substitutions in the two data sets differed by almost an order of magnitude (Table S2). Support for a rate matrix depends on genetic distance in the edge set. Better accuracy was associated with greater genetic distance. The lower accuracy of all algorithms on the 5-close data set for larger *k* is a plausible consequence of this effect. At the same time, the greater accuracy of IRM shows that it is better suited for optimal models selection than either HAL or ns-HAL. This lower accuracy of the HAL algorithms is expected given they are considering a larger possible solution space and thus has a greater chance for false positives.

The IRM was motivated by empirical evidence that tendencies for mutation are heritable. Such evidence manifests within a species where, for example, defects in mismatch DNA repair genes are associated with familial cases of colon cancer [Peltomäki, 2001]. This heritability of mutation tendencies also manifest between species. For instance, numerous species are known to have lost completely, or substantially, the capacity to methylate their DNA and these properties are shared among closely related lineages [Capuano et al., 2014]. IRM explicitly captures this tendency. The mincut algorithm improves the efficiency of tree traversal by evaluating longer edges first. If, as seems plausible, the likelihood of evolving distinctive mutation processes is proportional to chronological time then this approach should prove efficient. While it remains a possibility that lineages may independently evolve the exact same mutation tendencies, a prospect captured by HAL and not IRM, such convergent evolution seems less likely. Therefore, IRM serves as a valuable null hypothesis for examining whether lineages exhibiting comparable gene loss exhibit comparable substitution matrices, e.g. gene loss affecting 5mC.

Given its consistency with biological drivers underpinning mutation, we suggest fast-IRM is well suited as a model selection strategy for non-stationary process models. On the 50-taxon naked coral alignments with the second codon sites, we found that IRM and fast-IRM were consistent. In terms of compute speed, both fast-IRM and IRM were superior to ns-HAL for all values of *k*. The ∼ 2× speed-up of fast-IRM over IRM along with the identical inferences (Table 6) shows fast-IRM is to be preferred. We expect this performance difference will increase with increasing number of taxa.

## Supporting information

Supplementary tables and algorithms

## Supporting Information

Algorithms are described in more detail in the Supplementary file.

## Acknowledgements

We thank Teresa Neeman for suggesting the McNemar test and Helmut Simon for comments on an earlier version of the manuscript.

