## Supplementary tables and algorithms for "The Inherited Rate Matrix algorithm for phylogenetic model selection for non-stationary Markov processes"

### Algorithms for IRM

We present implementations of the core algorithms central to IRM below. These are presented as Python3 code. The code is not exactly identical to that in the distributed source code due to simplifications made for readability. Table S1 identifies the counterpart in `phylotree.irm`.

| **Below** | `phylotree.irm` |
|---|---|
| `variant_min_cut()` | `variant_min_cut()` |
| `get_edges()` | part of `get_edgeset()` |
| `cut_cluster_graph()` | part of `generate_cluster_graph()` and `cut_off_child()` |
| `cut_model_graphv` | `generate_cluster_graph()` |
| `is_cuttable()` | `is_cuttable()` |

Table S1: Mapping between functions below and the implementation in `phylotree.irm`

| | **5-close** | | **5-dispersed** | |
|---|---|---|---|---|
| ***k*** | **ns-HAL** | **IRM** | **ns-HAL** | **IRM** |
| **1** | 2081 | 2081 | 17923 | 17923 |
| **2** | 2099 | 2099 | 17840 | 17840 |
| **3** | 2091 | 2091 | 17780 | 17780 |
| **4** | 2091 | 2091 | 17700 | 17700 |
| **5** | 2081 | 2081 | 17784 | 17783 |
| **6** | 2079 | 2079 | 17751 | 17751 |
| **7** | 2097 | 2098 | 17750 | 17749 |

Table S2: Substitution number comparison between algorithm ns-HAL and algorithm IRM on 5-close and 5-dispersed species. Here, the values in table are round number of average ENS of 10 simulated alignments

**Algorithm S1** variant minimum cut algorithm

```
# mincut IRM model space generator
def variant_min_cut(tree):
    # Sort in descending order of edge/branch ENS distance
    ordered_edges = get_edges(tree, sort=True)
    num_edges = len(ordered_edges)
    # Graph stores all the parent models
    # Only a graph from the whole tree initially
    graphs = [[tree.deepcopy()]]
    start = 0
    while start < num_edges:
        # next_generation_graphs stores new cluster graphs
        # (all the children models) in the current generation
        next_generation_graphs = []
        stop = start + 1
        start_edge = ordered_edges[start]
        # Finding nodes with the same edge/branch length
        # as one of start edge
        for i in range(stop, num_edges):
            if start_edge.length != ordered_edges[i].length:
                break
            stop = i + 1

        # Cut each graph to generate a new graph with
        # more subgraphs
        for cluster_graph in graphs:
            # The corresponding 2-dimension edge set of
            # parent model
            parent_model_edge_set = \
                get_model_edgeset(cluster_graph)
            # The corresponding 3-dimension edge set of
            # children models generated by parent model
            children_models_edge_set = []
            for cut_id in range(start, stop):
                # Find the subgraph with cutting edge and cut
                # it into two smaller subgraphs to generate a
                # new model
                new_model_graph = cut_model_graph( \
                    cluster_graph, ordered_edges[cut_id].name)
                if new_model_graph:
                    new_edges = get_model_edgeset(new_model_graph)
                    children_models_edge_set.append(new_edges)
                    next_generation_graphs.append(new_model_graph)
            if children_models_edge_set:
                yield parent_model_edge_set, children_models_edge_set
            else:   # Not cut, pass to next generation
                next_generation_graphs.append(cluster_graph)

        graphs = next_generation_graphs
        start = start + 1
```

**Algorithm S2** Edge and graph selection utility functions

```
# Construct edge array from a tree
def get_edges(tree, sort=True):
    edges = [(node.length, node) for node in tree.preorder()
             if not node.is_root()]
    if sort:  # Make descending by edge/branch length
        edges.sort(reverse=True)
    return [e for _, e in edges]

# Construct model edgeset (a 2-dimension array)
def get_model_edgeset(cluster_graph, sort=True):
    edge_names_set = []
    for subgraph in cluster_graph:  # Traverse subgraph
        edges = get_edges(subgraph)
        edge_names = [edge.name for edge in edges]
        if not subgraph.is_root():
            edge_names.append(subgraph.name)
        if sort:
            edge_names.sort() # alphabetical sort
        edge_names_set.append(edge_names)
    if sort:
        edge_names_set.sort() # alphabetical sort
    return edge_names_set
```

**Algorithm S3** Graph and model cutting utility functions

```
def is_cuttable(node):
    if node.is_root():
        return False
    parent = node.parent
    return not parent.is_root() or len(parent.children) > 1

# graph is modified, so user must make sure they copy it
def cut_cluster_graph(graph, node_name):
    node = graph.get_node_matching_name(node_name)
    detached_child = node.parent.pop(node.index_in_parent())
    node.parent = None
    return node, graph

# cut a model graph
def cut_model_graph(model_graph, node_name):
    new_model_graph = []
    for subgraph in model_graph:
        new_subgraph = None
        # Check whether the subgraph includes the node
        # If yes, split the matched source subgraph into 2
        for node in subgraph.preorder():
            if node.name == node_name:
                if not is_cuttable(node):
                    return None # Unnecessary to cut

                # Construct a subgraph from the parent node
                copied = subgraph.copy_topology()
                new_subgraph, cut_subgraph = \
                    cut_cluster_graph(copied, node_name)
                # Add new subgraph at the beginning
                new_model_graph.insert(0, new_subgraph)
                # Add source subgraph which is cut
                new_model_graph.append(cut_subgraph)
                break
        if new_subgraph is None:
            new_model_graph.append(subgraph)
    return new_model_graph
```
